# Peptidoglycan precursor synthesis along the sidewall of pole-growing mycobacteria

**DOI:** 10.1101/292607

**Authors:** Alam García-Heredia, Amol Arunrao Pohane, Emily S. Melzer, Caleb R. Carr, Taylor J. Fiolek, Sarah R. Rundell, Hoong Chuin Lim, Jeffrey Wagner, Yasu S. Morita, Benjamin M. Swarts, Caleb R. Carr, M. Sloan Siegrist

## Abstract

d-amino acid probes label cell wall peptidoglycan at both the poles and sidewall of pole-growing mycobacteria. Since peptidoglycan assembly along the cell periphery could provide a rapid, growth-independent means by which to edit the cell wall, we sought to clarify the precise metabolic fates of these probes. d-amino acid monopeptides were incorporated into peptidoglycan by l,d-transpeptidase remodeling enzymes to varying extents. Dipeptides were incorporated into cytoplasmic precursors. While dipeptide-marked peptidoglycan synthesis at the poles was associated with cell elongation, synthesis along the periphery was highly responsive to cell wall damage. Our observations suggest a post-expansion role for peptidoglycan assembly along the mycobacterial sidewall and provide a conceptual framework for understanding cell wall robustness in the face of polar growth.

## Introduction

Model, rod-shaped organisms such as *Escherichia coli* and *Bacillus subtilis* elongate across a broad swath of the cell (1, 2). Mycobacterial cells, by contrast, extend from narrower polar regions (3–9). Circumscription of growth to discrete zones poses spatial challenges to the bacterial cell. For example, if polar growth and division are the only sites of cell wall synthesis in mycobacteria, the entire lateral surface of the cell must be inert (3, 10–12). Such an expanse of non-renewable surface could leave the cell vulnerable to environmental or immune insults.

Since cell wall peptidoglycan synthesis is critical for bacterial replication, it is often used to localize the sites of growth and division. Intriguingly, d-amino acid probes, which in other species have been shown to incorporate into peptidoglycan (1, 11, 13), label both the poles and sidewall of mycobacteria (5, 13–17). The localization of these molecules is supported by the detection of peptidoglycan synthetic enzymes at the mycobacterial cell tips and periphery (5, 9, 18–20). However, both intracellular and extracellular incorporation pathways have been characterized or hypothesized for d-amino acid probes, complicating the interpretation of labeling patterns (21). Intracellular uptake implies that the probe enters the biosynthetic pathway at an early stage, and therefore marks nascent cell wall. Extracellular incorporation, on the other hand, suggests that the probe enters the pathway at a later stage and/or is part of enzymatic remodeling of the macromolecule in question. The extent to which peptidoglycan synthesis and remodeling are linked is not clear (10, 22, 23) and may vary with species and external milieu. In *Mycobacterium tuberculosis*, for example, there is indirect but abundant data that suggest that there is substantial cell envelope remodeling during infection when growth and peptidoglycan synthesis are presumed to be slow or nonexistent (7).

An intracellular metabolic tagging method for the cell wall would be an ideal tool for determining whether tip-extending mycobacteria can synthesize peptidoglycan along their lateral surfaces. At least two pieces of evidence suggest that d-alanine-d-alanine dipeptide probes are incorporated into peptidoglycan via the cytoplasmic MurF ligase (24, 25). First, derivatives of d-alanine-d-alanine rescue the growth of *Chlamydia trachomatis* treated with d-cycloserine, an antibiotic that inhibits peptidoglycan synthesis by inhibiting the production and self-ligation of d-alanine in the cytoplasm (24). Second, *B. subtilis* cells stripped of mature peptidoglycan by lysozyme treatment retain a small amount of dipeptide-derived fluorescence (25). While these data are suggestive, formal demonstration of intracellular incorporation requires direct evidence that the probe labels peptidoglycan precursors. More broadly, better characterization of the metabolic fate of probes would increase the precision of conclusions that can be drawn from labeling experiments (26, 27).

Here we sought to delineate cell wall synthesis from remodeling in *Mycobacterium smegmatis* and *M. tuberculosis*. Monopeptide d-amino acid probes chiefly reported peptidoglycan remodeling by l,d-transpeptidases while dipeptides marked lipid-linked peptidoglycan precursors. All of the probes tested labeled the poles and sidewall of mycobacteria, indicating that cell wall metabolism in these regions comprises both synthetic and remodeling reactions. While peptidoglycan assembly along the mycobacterial periphery did not support obvious surface expansion, it was greatly enhanced by cell wall damage. Such activity may allow editing of a complex, essential structure at timescales faster than those permitted by polar growth.

## Results

### Metabolic labeling of mycobacterial envelope comprises asymmetric polar gradients

Mycobacteria have been shown to expand from their poles (3–9) but published micrographs suggest that d-amino acid probes may label both the poles and sidewall of these organisms (5, 13–17). Metabolic labeling can be achieved by a one-step process, in which the fluorophore is directly appended to the probe, or a two-step process in which a small chemical tag on the d-amino acid is detected by subsequent reaction with a fluorescent reactive partner ((21), **Figure 1A**). We first reexamined the localization of various d-amino acid probes reported in the literature, including RADA (11), which is directly conjugated to 5-carboxytetramethylrhodamine, and the two-step alkyne-d-alanine (alkDA or EDA, (11, 13)), and alkyne-d-alanine-d-alanine (alkDADA or EDA-DA, (24); we use the metabolic labeling nomenclature originally adopted in (28)) which we detected by copper-catalyzed azide-alkyne cycloaddition (CuAAC) after fixation (**Figures 1B, 1C**). After *M. smegmatis* cells were incubated for ~10% generation in probe, high resolution and quantitative microscopy revealed that they had asymmetric, bidirectional gradients of fluorescence that emanated from the poles and continued along the sidewall (**Figure 2A**). Polar gradients of dipeptide labeling were also apparent in live cells when we detected azido-d-alanine-d-alanine (azDADA or ADA-DA, (24)) incorporation by either CuAAC (using low copper, bio-friendly reaction conditions ((29), **Figure 2—figure supplement 1A**) or by copper-free, strain-promoted azide-alkyne cycloaddition (SPAAC, **Figure 2—figure supplement 1B**).

**Figure 1.**
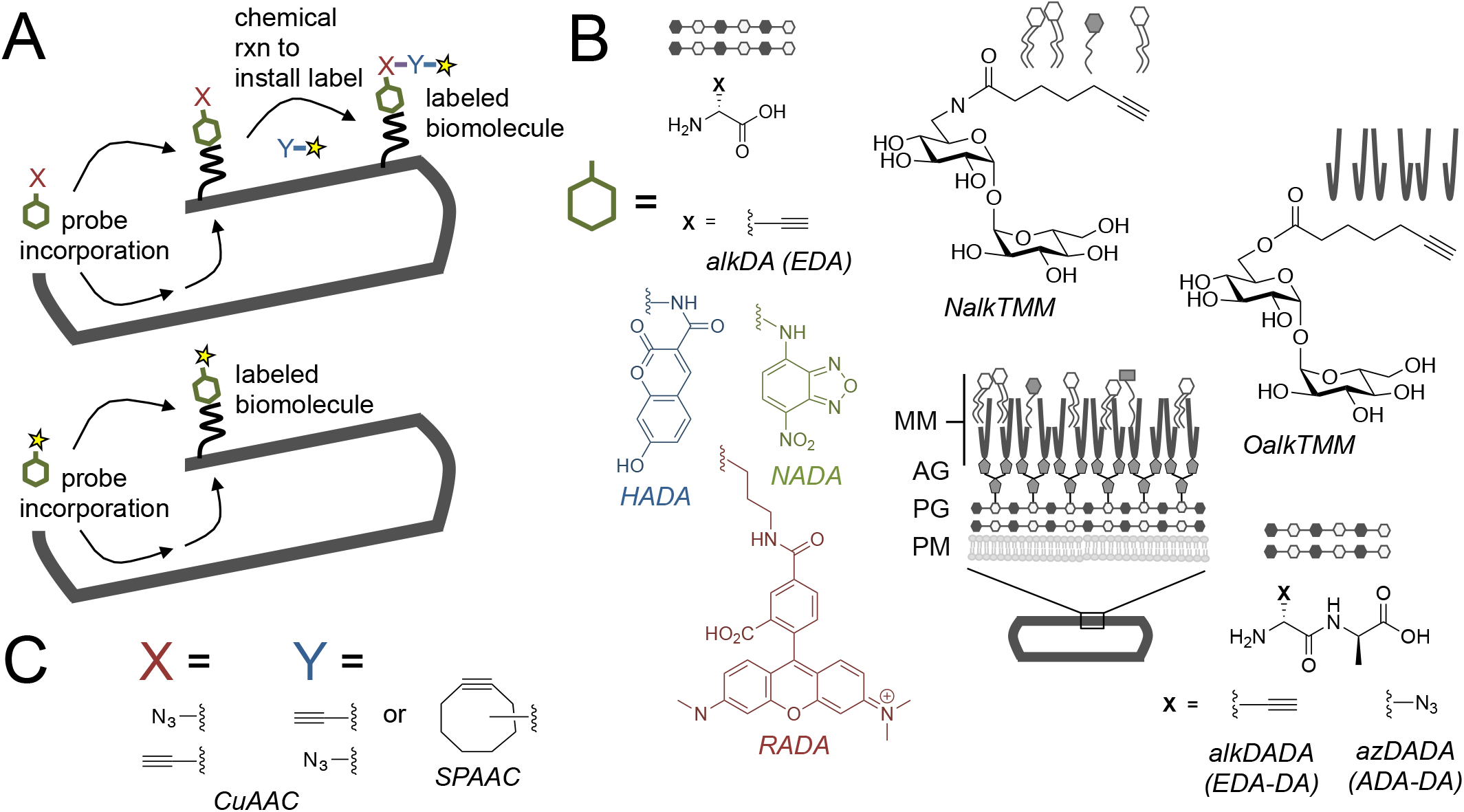
Cell envelope metabolic labeling in mycobacteria. A, Schematic of one and two step metabolic labeling. Top, a cell envelope precursor or ‘probe’ bearing a reactive group is incorporated into the envelope by the endogenous enzymatic machinery of the cell. The presence of the probe is then revealed by a chemical reaction with a label that bears a complementary reactive group. Bottom, in some cases the probe can be pre-labeled, bypassing the chemical ligation step and embedding the detection moiety directly into the macromolecule. Yellow star, fluorophore. See (21) for more details. B, Probes used in this work to mark the mycobacterial envelope. See text for details. Colored and black chemical structures denote probes used in one and two step labeling, respectively. MM, mycomembrane; AG, arabinogalactan; PG, peptidoglycan; PM, plasma membrane. C, X and Y reactive partners used in this work for two step labeling as shown in A. CuAAC, copper-catalyzed azide-alkyne cycloaddition; SPAAC, strain-promoted azide-alkyne cycloaddition.

**Figure 2.**
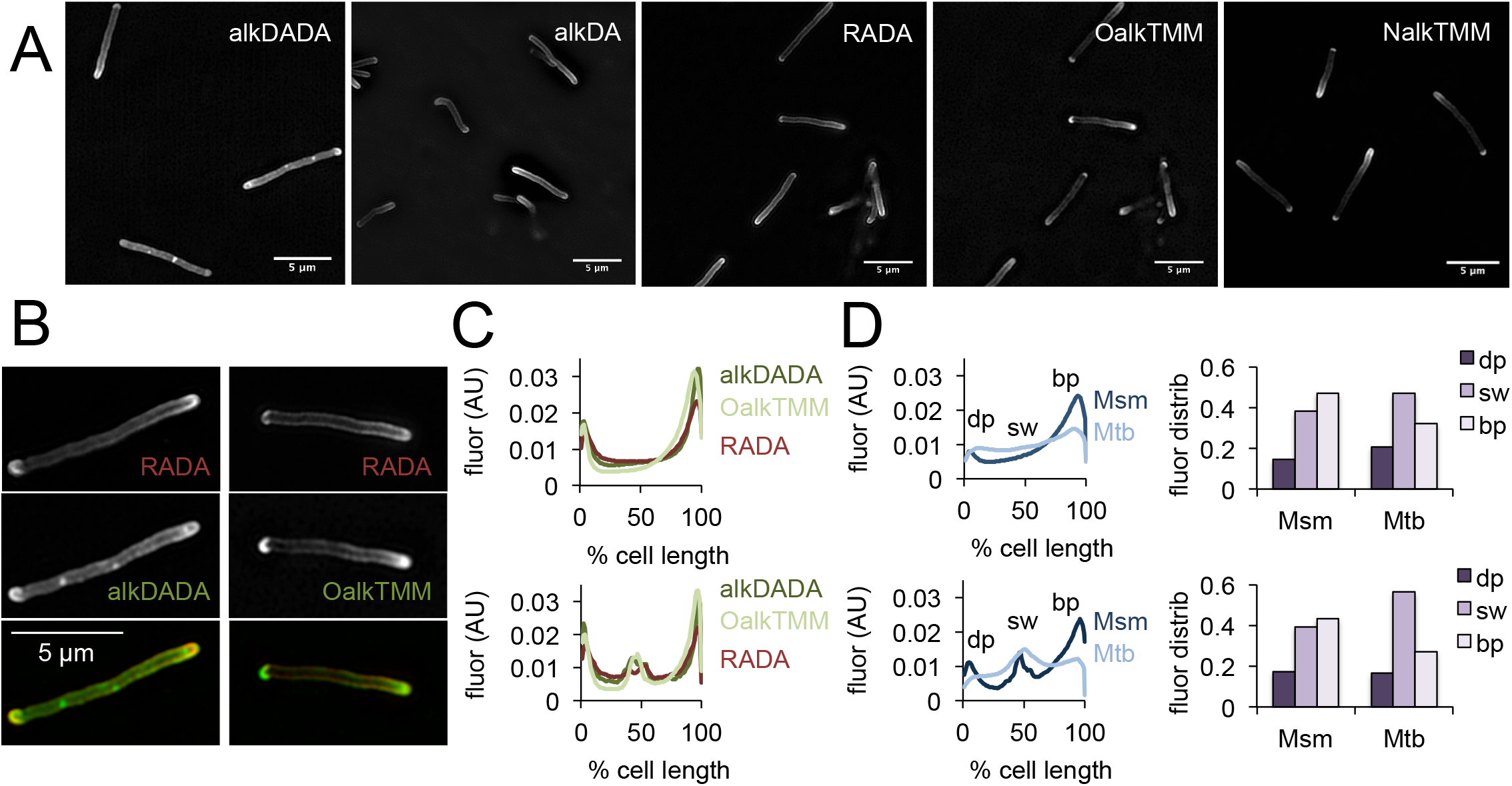
Asymmetric polar gradients of cell envelope metabolic labeling in mycobacteria. A, *M. smegmatis* was incubated for 15 min (~10% generation) in the indicated probe, then washed and fixed. Alkynyl probes were detected by CuAAC with azido-CR110 and cells were imaged by structured illumination microscopy. B, *M. smegmatis* dual labeled with RADA and alkDADA, left, or RADA and OalkTMM, right, and imaged by conventional fluorescence microscopy. C, *M. smegmatis* was labeled as in B and cellular fluorescence was quantitated for cells without (top; 77<n<85) or with (bottom; 9<n<51) visible septa for RADA, OalkTMM and alkDADA. Signal was normalized to cell length and to total fluorescence intensity. Cells were oriented such that the brighter pole is on the right hand side of the graph. D, *M. smegmatis* (Msm) and *M. tuberculosis* (Mtb) were labeled with HADA for 15 min and 2 hours, respectively, then washed and fixed. Fluorescence was quantitated as in C for cells without (top; 34<n<42) and with (bottom; 9<n<31) visible septa. We defined the dim pole (dp; dark purple) as the sum of the fluorescence intensity over the first 25% of the cell; the sidewall (sw; medium purple) as the sum from 25% to 75%; and the bright pole (bp; light purple) as the sum over the final 25% of the cell. Fluor distrib, fluorescence distribution. AU, arbitrary units.

The mycobacterial cell envelope is comprised of covalently-bound peptidoglycan, arabinogalactan and mycolic acids, as well as intercalated glycolipids and a thick capsule ((30), **Figure 1B**). Assembly of the envelope layers has long been presumed to be spatially coincident. This is largely based on biochemical data suggesting that ligation of arabinogalactan to peptidoglycan occurs concurrently with crosslinking of the latter by transpeptidases (31). We and others have found that cytoplasmic enzymes that mediate arabinogalactan and mycomembrane synthesis are enriched at the poles but also present along the periphery of the cell (5, 32, 33), as is metabolic labeling by OalkTMM and NalkTMM ((34–36), **Figure 2A**). OalkTMM and NalkTMM are trehalose monomycolate derivatives that predominantly mark covalent mycolates and trehalose dimycolate, respectively, in the mycomembrane (**Figure 1B**, (36)). Since the azido and alkynyl groups on the different probes are not orthogonal to each other (**Figure 1C**), we opted to compare peptidoglycan and mycomembrane labeling patterns by using RADA as a fiducial marker. The cell pole with brighter RADA fluorescence also had more alkDADA or OalkTMM labeling (**Figure 2B**), suggesting that the polar orientation of peptidoglycan and mycomembrane metabolism is coincident. We then compared the fluorescence intensity profiles of cells that had been individually labeled with the probes, and found similar, average distributions of RADA, alkDADA and OalkTMM at the poles and peripheries of the cells (**Figure 2C**).

We next sought to address whether the cell envelope of the related *M. tuberculosis* is also labeled in polar gradients. We previously showed that alkDA incorporates into the cell surface of the organism (13) but were unable to stain the entire population of bacteria. To investigate the origin of labeling heterogeneity, we first tested whether the structure of fluorophore (**Figure 1B**) influenced probe incorporation by incubating *M. tuberculosis* in HADA, NADA or RADA and assessing population fluorescence by flow cytometry. HADA and NADA incubation yielded well-defined fluorescent populations (**Figure 2—figure supplement 2A**). RADA also labeled the entire *M. tuberculosis* population, albeit with greater cell-to-cell variability in fluorescence intensity. Given that *M. tuberculosis* incorporates fluorescent probes HADA and NADA relatively evenly across the population, we hypothesized that the apparent heterogeneity that we previously observed for alkDA labeling (13) was the result of an inefficient CuAAC ligation. We obtained modest improvements by changing the reaction conditions, more specifically, by swapping the BTTP ligand (29, 37) for the TBTA ligand, altering the ratio of ligand:Cu(I) and increasing the azide label concentration. We also switched our detection moiety to an azide appended to hydroxycoumarin, the same small, uncharged fluorophore as the one-step HADA probe (**Figure 1B**). Under our optimized conditions we detected azido-coumarin fluorescence from ~15-20% of cells that had been incubated in alkDA, alkDADA or OalkTMM (**Figure 2— figure supplements 2B, 2C**).

Although unable to achieve homogenous *M. tuberculosis* labeling with two-step envelope probes, we decided to test whether the sites of envelope labeling in the limited fluorescent subpopulation resemble those of *M. smegmatis*. HADA, alkDADA and OalkTMM tagging all produced cells that had a mixture of sidewall and polar fluorescence (**Figure 2—figure supplement 2C**) but exhibited a higher degree of cell-to-cell variability compared to *M. smegmatis*. Quantitation of HADA fluorescence showed that peptidoglycan metabolism comprised asymmetric polar gradients when averaged across the population (**Figure 2D**). We next asked whether there was a cell-wide difference in labeling distribution in *M. tuberculosis* compared to *M. smegmatis*. We arbitrarily defined the dimmer polar region as the first 25% of the cell length, the sidewall as the middle 50%, and the brighter polar region as the final 25%. In a cell with perfect, circumferential labeling, the ratio of dim pole:sidewall:fast pole in a cell would be 25:50:25. Since HADA labeled a large proportion of *M. tuberculosis* (**Figure 2—figure supplement 2A**) and was more resistant to photobleaching than NADA, we compared these ratios for HADA in septating and non-septating *M. smegmatis* and *M. tuberculosis* labeled for ~10% generation time (**Figure 2D**). For *M. smegmatis*, the ratios were 15:38:47 for non-septating cells and 17:39:43 for septating cells, while for *M. tuberculosis*, they were 21:47:32 and 17:56:27 for *M. tuberculosis*. These data suggest that a greater proportion of peptidoglycan labeling in *M. tuberculosis* likely occurs along the cell periphery than in *M. smegmatis*, although we cannot rule out a differential contribution of cyan autofluorescence in the two species (38).

### Intracellular and extracellular pathways of d-amino acid probe incorporation in mycobacteria

Since the cell periphery is not known to support surface expansion in mycobacteria (3–9), we sought to characterize the molecular processes that underlie d-amino acid labeling patterns. OalkTMM and NalkTMM are inserted directly by the extracellular Antigen 85 complex into the mycomembrane (36). However, there are three potential pathways by which d-amino acid probes might incorporate into mycobacterial peptidoglycan (21, 39, 40): an intracellular, biosynthetic route and two extracellular routes, mediated by d,d-transpeptidase or l,d-transpeptidase remodeling enzymes (**Figure 3A**). Peptidoglycan remodeling, particularly by the l,d-transpeptidases abundantly encoded in the mycobacterial genome, may not strictly correlate with synthesis of the biopolymer (10, 11, 22, 23). Therefore, we sought to distinguish the different routes of incorporation for d-amino acid probes in mycobacteria.

**Figure 3.**
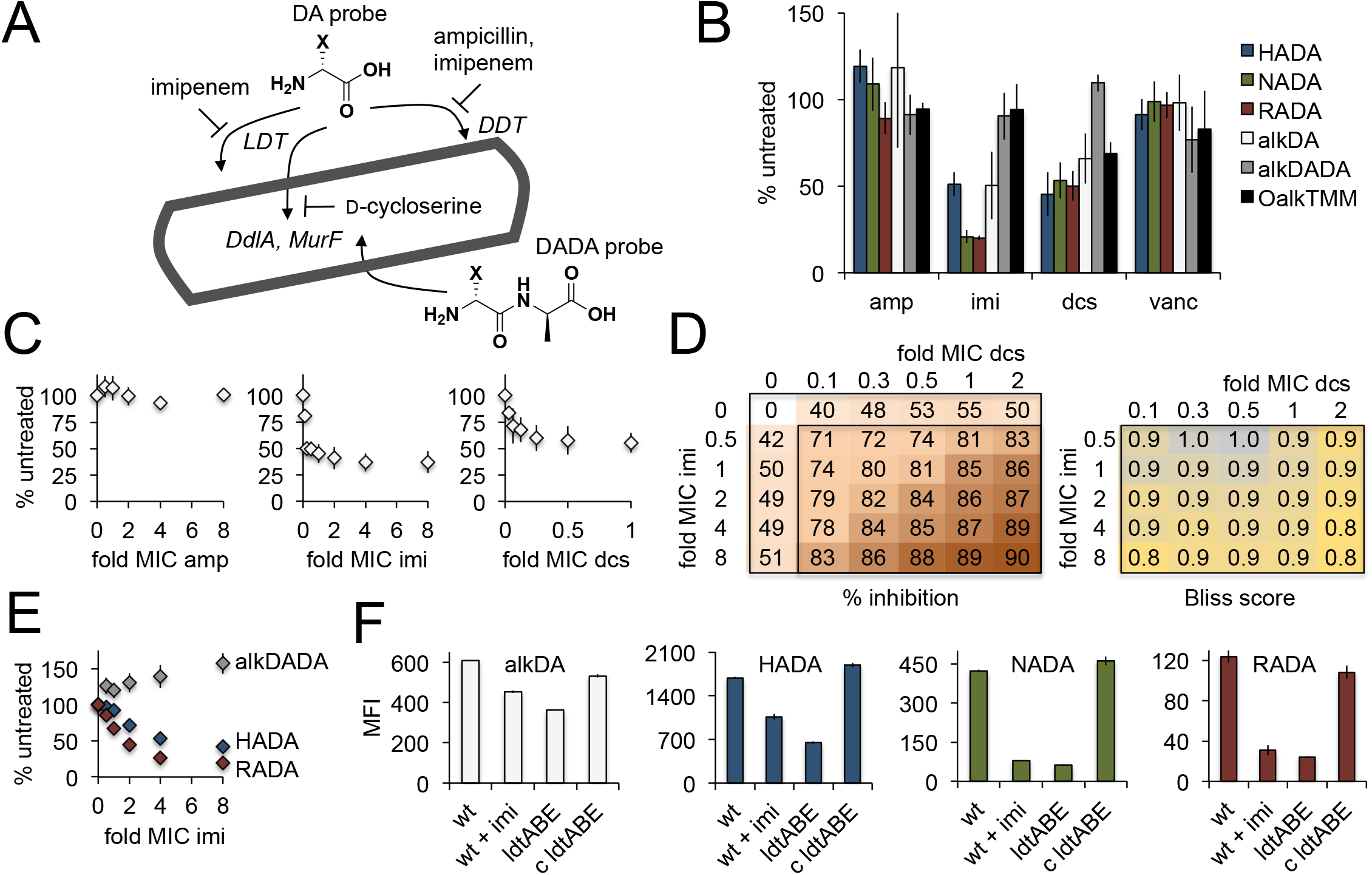
Multiple pathways of d-amino acid probe incorporation in *M. smegmatis*. A, Schematic of the theoretical routes of d-amino acid (DA) and d-alanine d-alanine (DADA) probe incorporation. LDT, l,d-transeptidase, DDT, d,d-transpeptidase (DDTs). For more details on the peptidoglycan synthesis pathway, see **Figure 3—figure supplement 1**. B, Sensitivity of HADA (blue), NADA (green), RADA (red), alkDA (light grey), alkDADA (dark grey) and OalkTMM (black) to antibiotics. Imi, imipenem + clavulanate; amp, ampicillin + clavulanate; dcs, d-cycloserine; vanc, vancomycin. *M. smegmatis* was pretreated or not with the indicated antibiotics at 2X MIC for 30 min then incubated an additional 15 min in the presence of probe. The bacteria were then washed and fixed. The alkyne-bearing probes were detected by CuAAC with azido-CR110 and quantitated by flow cytometry. Experiment was performed 3-4 times in triplicate. For each biological replicate, the averaged median fluorescence intensities (MFI) of the drug-treated samples were divided by the MFI of untreated bacteria. Data are expressed as the average percentage of untreated labeling across the biological replicates. Error bars, +/- standard deviation. C, Effect of antibiotic dose on alkDA-derived fluorescence. *M. smegmatis* was pretreated or not with drugs at the fold-MIC indicated and labeled as in B. Experiment was performed 3 times in triplicate. For each biological replicate, the averaged MFI of the control (no drug, no alkDA but subjected to CuAAC) was subtracted from the averaged MFI of the drug-treated sample. This was then divided by the averaged MFI of untreated control (no drug but incubated in alkDA and subjected to CuAAC) from which the control MFI had also been subtracted. Data are expressed as the average percentage of untreated labeling across the biological replicates. Error bars, +/- standard deviation. D, Left, combined effects of imipenem and d-cycloserine on alkDA-derived fluorescence. *M. smegmatis* was pretreated or not with the drugs at the fold-MIC indicated and labeled as in B. Experiment was performed twice in triplicate with similar results. One data set is shown. The percent of untreated labeling was calculated as in C and subtracted from 100 to obtain the percent inhibition. Right, Bliss interaction scores for each pair of doses in left-hand graph were calculated as (E_I_+E_D_-E_I_E_D_)/E_I,D_ where E_I_ is the effect of imipenem at dose *i*, E_D_ is the effect of d-cycloserine at dose *d* and E_I,D_ is the observed effect of the drugs at dose *i* and dose *d*. Combinations that produce Bliss scores greater than, equal to, or less than 1 are respectively interpreted as antagonistic, additive, or synergistic interactions. E, Dose-dependent effect of imipenem on alkDADA (grey), HADA (blue) and RADA (red) labeling. *M. smegmatis* was pretreated or not with imipenem at the fold-MIC indicated and labeled as in B. Experiment was performed 3 times in triplicate and one representative data set is shown. For each technical replicate, the averaged median fluorescence intensities (MFI) of the drug-treated samples were divided by the averaged MFI of untreated bacteria. Data are expressed as the average percentage of untreated labeling across the technical replicates. Error bars, +/- standard deviation. F, Wildtype *M. smegmatis* pre-treated or not with 2X MIC imipenem and untreated *ΔldtABE* and complement (c ldtABE) were labeled with alkDA (white), HADA (blue), NADA (green) or RADA (red) and processed as in B. Experiment was performed 2-10 times in triplicate. Representative data from one of the biological replicates is shown here. Error bars, +/- standard deviation.

It seemed possible that the chemical structure of the derivative (**Figure 1B**) and/or number of labeling steps (**Figure 1A**) might influence probe uptake, so we first tested the labeling sensitivity of a panel of d-amino acid derivatives to antibiotics that inhibit potential incorporation routes (**Figure 3A**). We also assessed OalkTMM and the d-alanine-d-alanine dipeptide probe, the latter of which has been proposed to tag the peptidoglycan of other species via the cytoplasmic MurF ligase (**Figures 3A, figure supplement 1**, (24, 25)). d-cycloserine is a cyclic analog of d-alanine that inhibits the d-alanine racemase (Alr) and ligase (DdlA) in mycobacteria (41). Together with the β-lactamase inhibitor clavulanate, β-lactams like ampicillin block d,d-transpeptidases and d,d-carboxypeptidases. Broader-spectrum carbapenems such as imipenem additionally inhibit l,d-transpeptidases (42). We also included vancomycin, an antibiotic that interferes with transpeptidation and transglycosylation by steric occlusion, to control for general defects in periplasmic peptidoglycan assembly. We empirically determined a time frame of drug treatment that did not compromise *M. smegmatis* viability (**Figure 3—figure supplement 2**). Within this time frame, labeling for all single residue d-amino acid probes decreased significantly in response to imipenem or d-cycloserine treatment (**Figure 3B**). By contrast, labeling by the dipeptide probe was relatively resistant to these antibiotics. Neither vancomycin nor ampicillin had a clear effect on labeling by any of the probes, indicating that cell death, disruption of peptidoglycan polymerization, and abrogration of d,d-transpeptidation do not explain the metabolic incorporation differences. OalkTMM labeling was sensitive to d-cycloserine, but not to imipenem nor ampicillin, suggesting that transfer of mycolates to arabinogalactan may require peptidoglycan precursor synthesis but not transpeptidation.

To test whether distinct mechanisms of probe incorporation were inhibited by d-cycloserine and imipenem, we performed a chemical epistasis experiment. We first examined the effect of different drug concentrations on alkDA incorporation (**Figure 3C**). Treatment with either d-cycloserine or imipenem, but not ampicillin, resulted in dose-dependent inhibition of the probe-derived fluorescence that plateaued at approximately half of untreated levels. We then assessed the combined effect of d-cycloserine and imipenem by Bliss independence (43), a commonly used reference model for predicting drug-drug interactions based on the dose response for the individual drugs. This method is most appropriate when dual inhibition proceeds via distinct mechanisms *e.g*. activity against different molecules, enzymes or pathways (44). The effects of d-cycloserine and imipenem on alkDA incorporation were the same or slightly greater than the potencies predicted by the antibiotics individually (**Figure 3D**). The primarily-additive nature of these antibiotics in the Bliss independence model is consistent with the idea that d-cycloserine and imipenem block distinct pathways of alkDA incorporation, and therefore that the probe incorporates into mycobacterial peptidoglycan via both cytoplasmic and l,d-transpeptidase routes.

RADA and NADA labeling were more sensitive to imipenem than HADA and alkDA (**Figure 3B**), at multiple concentrations of drug (**Figure 3E**). To test whether the probes were differentially incorporated by l,d-transpeptidases, we knocked out three of the six enzymes encoded in the *M. smegmatis* genome. We chose to focus on LdtA, LdtB and LdtE since the *M. tuberculosis* homologs (Ldt_MT1_, Ldt_MT2_, and Ldt_MT4_, respectively) have been shown *in vitro* to have both cross linking and d-amino acid exchange activity and to be inhibited by imipenem (45). RADA and NADA fluorescence decreased by ~80-85% the absence of *ldtABE*, and this effect was complemented by the expression of *ldtA* alone (**Figure 3F**). We observed a more moderate effect on HADA and alkDA labeling, which were decreased by ~40-60%. Collectively these data suggest that: 1. RADA and NADA probes primarily report l,d-transpeptidase activity, tetrapeptide substrate, or both, and 2. HADA and alkDA likely label via both cytoplasmic and l,d-transpeptidase routes.

### Dipeptide d-amino acid probe incorporates into mycobacterial peptidoglycan precursors

Since RADA labeling in mycobacteria is largely indicative of l,d-transpeptidase activity (**Figure 3**) and occurs at the poles and along the sidewall (**Figure 2**), we surmise that peptidoglycan is remodeled at both of these locations. We wished to determine whether remodeling was coincident with biopolymer synthesis but were limited by the multiple incorporation routes of alkDA and HADA (**Figure 3**). Dipeptide d-amino acid probes have been proposed to report peptidoglycan synthesis in other species via a cytoplasmic, MurF-dependent pathway (**Figures 3A, figure supplement 1**, (24, 25)). Consistent with this notion, and in contrast to the monopeptide probes, alkDADA labeling was relatively stable to imipenem treatment (Figures 3B, 3E) and inefficiently incorporated (**Figure 4—supplement 1**). Thus we were surprised to observe that overall labeling by alkDADA decreased in the absence of LdtA, LdtB, LdtE or combinations thereof (Figure 4—supplement 2A). The reduction in signal occurred primarily at the poles (**Figure 4—supplement 2B**) yet loss of the enzymes did not impair bacterial growth (**Figure 4—supplement 3**). As we do not yet understand the mechanistic basis for this observation, we sought to directly test the hypothesis that dipeptide probes incorporate into peptidoglycan precursors.

d-alanine is produced in mycobacteria by Alr, the d-alanine racemase (**Figure 3—figure supplement 1**). The molecule is linked to a second d-alanine by DdlA, the d-alanine ligase, and the resulting dipeptide is added to the UDP-MurNAc-tripeptide by MurF. If the alkDADA probe is able to label mycobacterial peptidoglycan via MurF, addition of the molecule to the growth medium should rescue a mutant that is unable to make d-alanine-d-alanine. We first constructed an *alr* deletion mutant in *M. smegmatis* and verified that growth is rescued by exogenous d-alanine but not by alkDA (**Figure 4A**). Although our antibiotic data suggest that alkDA is incorporated into peptidoglycan in part via a cytoplasmic pathway (**Figure 3**), the inability of this probe to rescue growth was not surprising given the substrate specificities of Ddl and MurF (46) and inefficient synthesis of UDP-MurNAc-pentapeptide with d-amino acids other than alanine (39). In contrast to the alkDA results, we were able to rescue *alr* with either d-alanine-d-alanine or its alkynyl derivative (**Figure 4A**).

**Figure 4.**
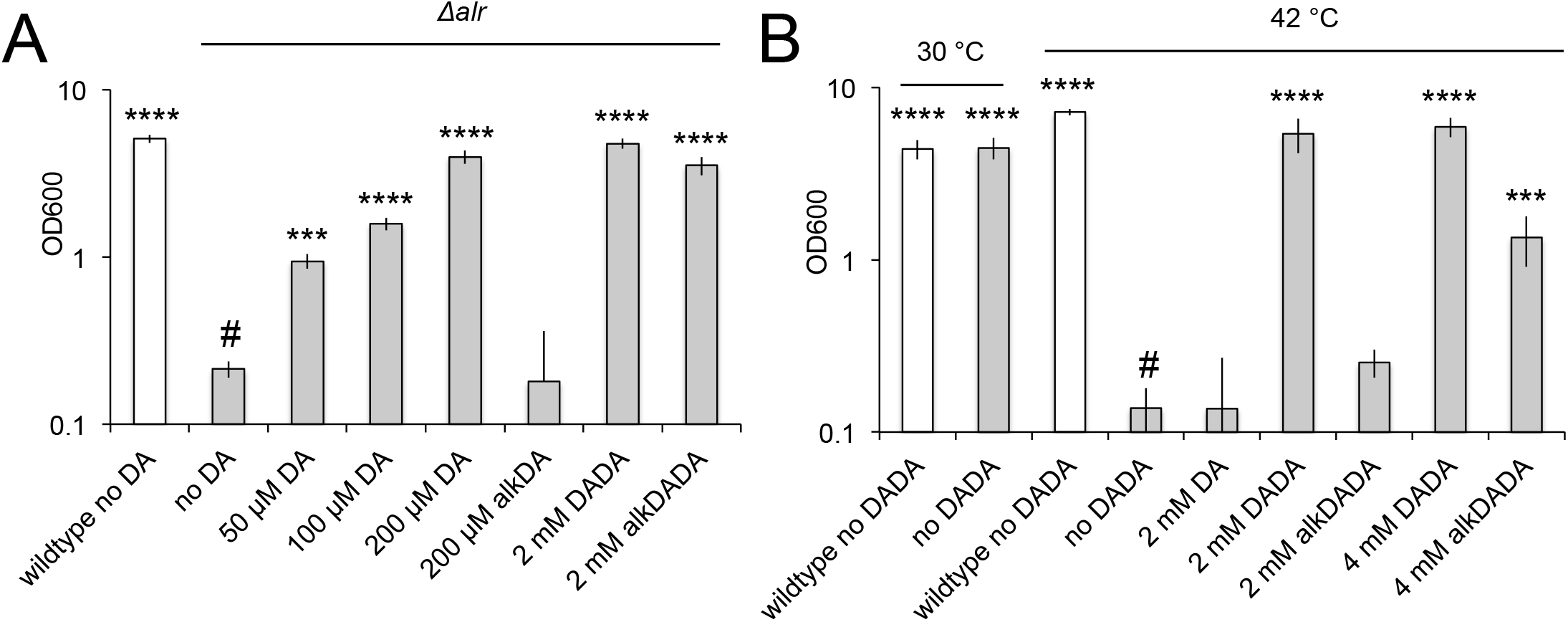
alkDADA rescues the growth of d-alanine racemase (Alr) and ligase (DdlA) mutants. alkDADA probe supports the growth of *Δalr*, A, or temperature-sensitive *ddlA*^ts^, B. Wildtype (white bars) and *Δalr* (grey bars), B, or *ddlA*^ts^ (grey bars), were grown in the presence or absence of exogenous d-alanine, d-alanine-d-alanine or alkynyl derivatives thereof. Error bars, +/- standard deviation. Significant differences compared to *Δalr* no d-alanine (#, second bar from left), A, or *ddlA^ts^* at 42 °C no d-alanine-d-alanine (#, fourth bar from left), B, One-way ANOVA with Dunnett’s test, are shown for 3-4 biological replicates. ***, p < 0.005, ****, p<0.0005.

We considered the possibility that alkDADA may be digested by a d,d-carboxypeptidase prior to incorporation. This would result in the release of both unlabeled d-alanine and alkDA, the first of which could account for *alr* growth rescue (**Figure 4A**). We reasoned that a more precise gauge for MurF-dependent incorporation of the intact probe would be whether it could support the replication of a strain unable to ligate d-alanine to itself. Therefore, we confirmed that alkDADA rescues the growth of a temperature-sensitive *ddlA* mutant (47) at the non-permissive temperature (**Figure 4B**).

Our genetic data supported a MurF-dependent pathway of alkDADA incorporation into peptidoglycan. If true, the probe should be present in precursors such as lipid I and lipid II (**Figure 3—figure supplement 1**). To test this hypothesis, we first optimized for mycobacteria a recently-reported protocol for detecting lipid-linked precursors (48, 49). We extracted lipidic species from *M. smegmatis* and exchanged endogenous d-alanines for biotin-d-lysine (BDL) *in vitro* using purified *Staphylococcus aureus* enzyme PBP4, a promiscuous d,d-transpeptidase (48). Biotinylated species were separated by SDS-PAGE and detected by horseradish peroxidase-conjugated streptavidin. We detected a biotin-linked, low molecular weight band that is reduced upon d-cycloserine treatment and accumulates when the lipid II flippase MurJ (MviN, (50)) is depleted or vancomycin is added (**Figure 5A**), conditions that have been shown to dramatically enhance precursor detection in other species (48, 49). These data strongly suggest that the BDL-marked species are lipid-linked peptidoglycan precursors. We next turned our attention to detecting lipid I/II from *M. smegmatis* incubated with alkDADA. Initially we were unable to identify alkDADA-labeled species from organic extracts of wildtype *M. smegmatis* that had been subjected to CuAAC ligation with picolyl azide biotin (**Figure 5B**). We reasoned that the proportion of labeled precursors might be below our limit of detection. Accordingly, we repeated the experiment in the *Δalr* background and found that we could clearly detect a low molecular-weight species band that accumulated with vancomycin treatment (**Figure 5B**, top) and that ran at the same size as BDL-labeled material (**Figure 5B**, bottom). We were also able to identify a low molecular weight species from *Δalr* incubated with azDADA that was revealed by either CuAAC or SPAAC ligation to alkyne- or cyclooctyne-biotin, respectively (**Figure 5C**). Taken together, our genetic and biochemical experiments show that the alkDADA and azDADA probes insert into mycobacterial peptidoglycan precursors by a MurF-dependent route.

**Figure 5.**
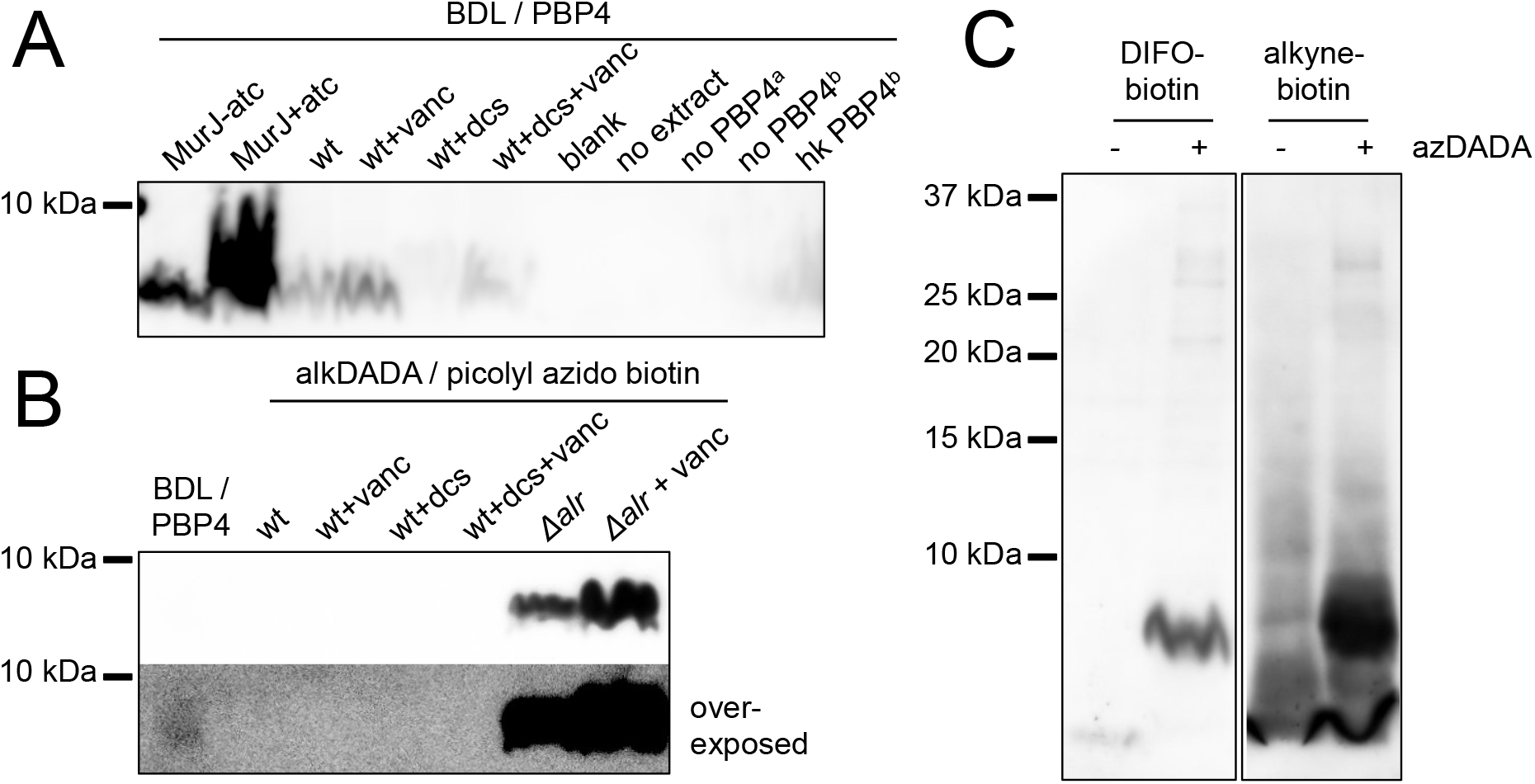
alkDADA and azDADA incorporate into lipid-linked peptidoglycan precursors. A, Detection of lipid-linked peptidoglycan precursors from organic extracts of *M. smegmatis*. Endogenous d-alanines were exchanged for biotin-d-lysine (BDL) via purified *S. aureus* PBP4. Biotinylated species detected by blotting with streptavidin-HRP. MurJ (MviN) depletion strain was incubated in anhydrotetracycline (atc) to induce protein degradation. Other strains were treated with vancomycin (vanc), d-cycloserine (dcs) or a combination prior to harvesting. Wt, wildtype; blank, no sample run; no extract, BDL and PBP4 alone ^a^, organic extract from MurJ-atc; ^b^, organic extract from MurJ+atc; hk, heat-killed. B, Detection of lipid-linked peptidoglycan precursors labeled by alkDADA *in vivo*. Wildtype and *Δalr* strains were incubated in alkDADA and treated or not with the indicated antibiotics prior to harvest. Alkyne-tagged species from organic extracts were ligated to picolyl azide biotin via CuAAC then detected as in A. BDL/PBP4, endogenous precursors from MurJ+atc were subjected to *in vitro* exchange reaction as in A. C, Detection of lipid-linked peptidoglycan precursors labeled by azDADA *in vivo. Δalr was* incubated in azDADA. Azide-tagged species from organic extracts were ligated to DIFO-biotin via SPAAC or to alkyne biotin via CuAAC then detected as in A.

### Fluorescent vancomycin and penicillin-binding proteins localize to the poles and sidewall in mycobacteria

The final, lipid-linked peptidoglycan precursor lipid II is synthesized by MurG on the cytoplasmic side of the plasma membrane then flipped to the periplasm and polymerized (**Figure 3—figure supplement 1**, (51)). We previously showed that MurG fused to two different fluorescent proteins and expressed under two different promoters is present at both the poles and periphery of *M. smegmatis* (5). Labeling by alkDADA marks similar subcellular locations even with pulses as short as ~1% generation time (**Figure 5—figure supplement 1**). These data suggest that lipid-linked peptidoglycan precursors are synthesized at lateral sites in addition to their expected localization at the poles. However, our standard experimental protocol for detecting envelope labeling is to perform CuAAC on fixed cells. Because formaldehyde fixation can permeabilize the plasma membrane to small molecules, labeled material may be intracellular, extracellular or both. Dipeptide labeling could therefore read out lipid I/II on the cytoplasmic face of the plasma membrane, uncrosslinked lipid II on the periplasmic side, or polymerized peptidoglycan.

To shed light on the potential fate(s) of peptidoglycan precursors made at different subcellular sites, we first stained live mycobacterial cells with fluorescent vancomycin. This reagent binds uncrosslinked peptidoglycan pentapeptides and does not normally cross the plasma membrane. Pentapeptide monomers are a low abundant species in *M. tuberculosis, M. abscessus* and *M. leprae* peptidoglycan (42, 52–54), suggesting that fluorescent vancomycin primarily reports extracellular, lipid-linked precursors in this genus. Labeling of *M. smegmatis* with this probe revealed both revealed both polar and lateral patches (**Figure 5—figure supplement 2**) as previously noted (8). This observation suggests that at least some of the peptidoglycan precursors present along the periphery of the mycobacterial cell are flipped to the periplasm.

We next sought to address whether these molecules could be used to build the peptidoglycan polymer. Transglycosylases from both the PBP (penicillin-binding proteins) and SEDS (shape, elongation, division, and sporulation) families stitch peptidoglycan precursors into the existing meshwork (**Figure 3—figure supplement 1**, (14, 32, 51, 55–58)). If peptidoglycan precursors are polymerized along the lateral surface of the mycobacterial cell, at least a subset of these periplasmic enzymes must be present at the sidewall to assemble the biopolymer. Two conserved PBPs in mycobacteria are likely responsible for most of the peptidoglycan polymerization required for cell viability, PonA1 and PonA2 (7, 19, 59). Published images of PonA1-mRFP and PonA1-mCherry localization suggested that the fusion proteins might decorate the mycobacterial sidewall in addition to the cell tips (9, 18, 19), but the resolution of the micrographs did not allow for definitive assignment. Therefore, we first verified the localization of PonA1-mRFP. We found that a subset of this fusion protein indeed homes to the lateral cell surface (**Figure 5—figure supplement 3A**).

We were concerned that overexpression of PonA1-mRFP causes aberrant polar morphology and is toxic to *M. smegmatis* (18, 19) and about the propensity of mCherry to cluster (60). Since our attempts to produce PonA1 fusions with different fluorescent proteins were unsuccessful, we opted to take a complementary, activity-based approach. Fluorescent derivatives of β-lactam antibiotics bind specifically and covalently to PBPs, and therefore have been used to image active enzyme in both protein gels and intact cells (61). Our images of whole cells labeled with Bocillin, a BODIPY conjugate of penicillin, were in agreement with those from a previous publication (20), and seemed to indicate that Bocillin binds both the poles and sidewall of *M. smegmatis* (**Figure 5—figure supplements 3B, 3C**). However, given the hydrophobicity of the BODIPY dye, we considered the possibility that Bocillin might nonspecifically associate with the greasy mycomembrane. Fluorescence across the cell surface was diminished by pre-treating cells with the β-lactam ampicillin, which prevents peptidoglycan assembly by binding to PBPs, but not d-cycloserine, which inhibits peptidoglycan synthesis in a PBP-independent manner (**Figure 5—figure supplements 3B, 3C**). These experiments suggest that at least some of the sidewall labeling of Bocillin is specific, and therefore, that PBPs are present and active in these locations.

### Expansion of the mycobacterial envelope is concentrated at the poles

Our data indicate that peptidoglycan precursors are made and likely polymerized both at the poles and sidewall. Peptidoglycan synthesis is often presumed to mark sites of bacterial cell growth. However, dispersed elongation has not been reported in mycobacteria. Accordingly we performed a pulse chase experiment to test whether cell expansion correlates with sites of metabolic labeling. After marking peptidoglycan with RADA, we tracked labeled and unlabeled cell surface during 15 min (~10% generation time) of outgrowth. While we cannot rule out sidewall expansion below our limit of detection, the fluorescence dilution in this experiment was consistent with previous reports ((3–5, 7, 14, 62) and restricted to the mycobacterial poles (**Figure 5—figure supplement 4**).

### Muramidase treatment increases peptidoglycan synthesis along the sidewall

What is the function of peptidoglycan assembly that does not directly contribute to physical expansion of the cell? We hypothesized that one role of growth-independent cell wall synthesis might be repair. More specifically, we reasoned that insertion of peptidoglycan building blocks directly along the cell periphery would enable a real-time, comprehensive response to damage (**Figure 6A**). Cell wall repair that is restricted to sites of mycobacterial growth, by contrast, would be confined to the poles and renew the cell surface only after several generations. Extended incubation of *M. smegmatis* (~48 hours) with the peptidoglycan-degrading enzyme lysozyme substantially decreases colony-forming units (63). We have also shown that spheroplasts generated by combined glycine and lysozyme treatment lack peptidoglycan (64). Together these data indicate that the enzyme is able to access and damage peptidoglycan in intact cells. We challenged *M. smegmatis* for 30 minutes in a mixture of lysozyme and mutanolysin, another enzyme that has been extensively used for *in vitro* digestion of peptidoglycan. After washing away the enzyme we assessed the sites of peptidoglycan synthesis by alkDADA labeling. Pre-treatment by the muramidases clearly shifted the fluorescence from the brighter pole towards the sidewall (**Figure 6B**). These data indicate that mycobacteria reallocate peptidoglycan assembly away from the faster-growing pole and towards the periphery upon damage to the cell wall (**Figure 6A**).

**Figure 6.**
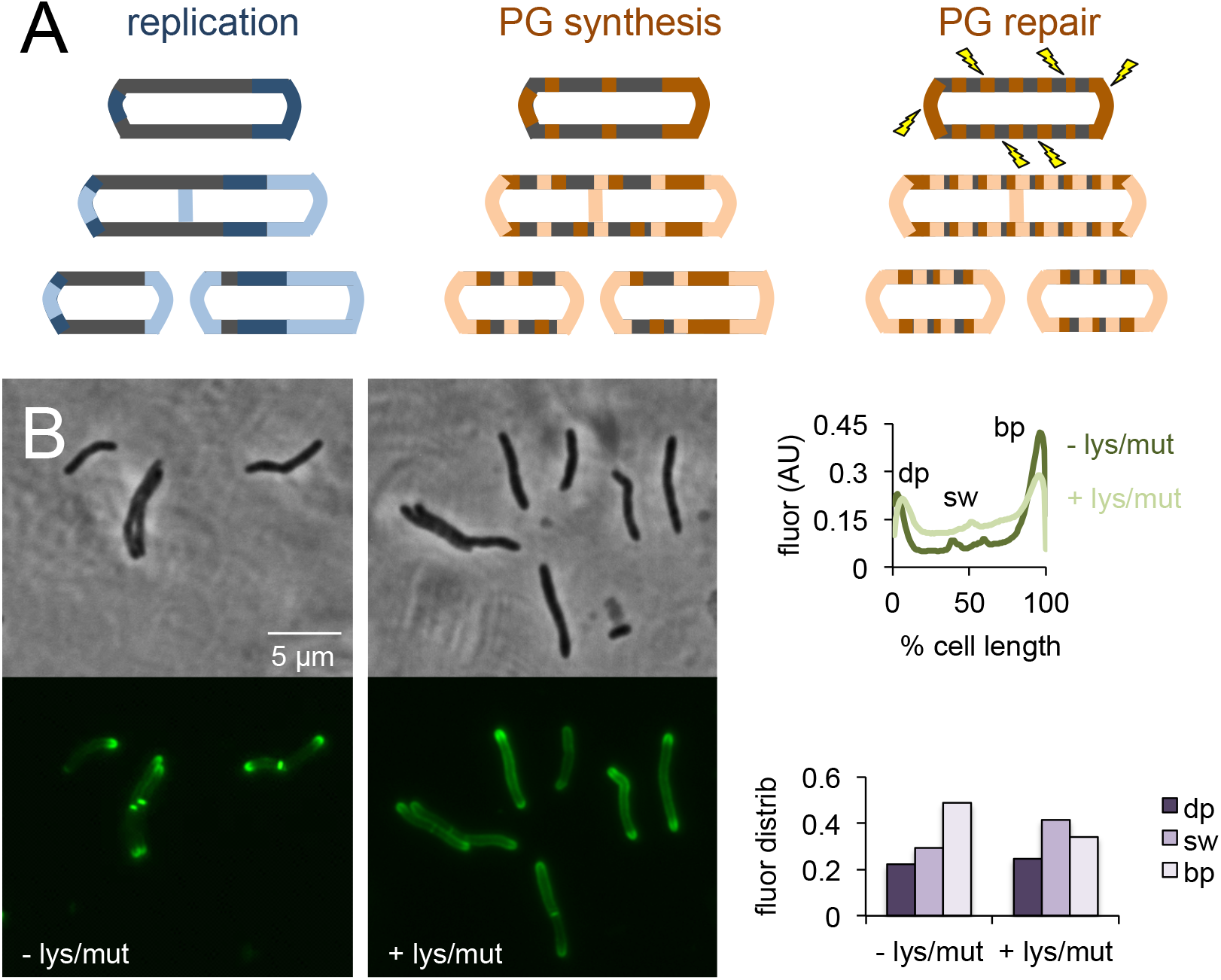
Peptidoglycan synthesis is redistributed to the sidewall upon cell wall damage. A, Model for the spatial organization of peptidoglycan (PG) synthesis and repair with respect to mycobacterial growth and division. Left, regions of cell surface that expand are highlighted in blue. Middle and right, areas of peptidoglycan precursor synthesis are highlighted in orange. B, *M. smegmatis* was pretreated or not with lysozyme and mutanolysin for 30 min then incubated an additional 15 min in the presence of alkDADA. The bacteria were then washed and fixed and subjected to CuAAC with azido-CR110. Images obtained by conventional fluorescence microscopy were quantified for 25<n<40 cells as in **Figure 2** except that i. signal was not normalized to total fluorescence intensity, only to cell length, and ii. cells with and without visible septa were analyzed together. Experiment was performed twice with similar results. dp, dim pole (dark purple); sw, sidewall (medium purple); bp, bright pole (light purple). Fluor distrib, fluorescence distribution. AU, arbitrary units.

## Discussion

In this work we aimed to address the seemingly discrepant observations that, on the one hand, mycobacteria expand from their tips (**Figure 5—figure supplement 4**, (3–5, 7, 14)), and on the other, metabolically-labeled cell wall and synthetic enzymes are detectable at both the poles and along the sidewall (**Figures 2, 5—figure supplement 3**, (5, 9, 18–20)). The first step to resolving this conundrum was to unambiguously identify sites of peptidoglycan synthesis. Although the d-amino acid probes that we and others have developed for peptidoglycan labeling have been extensively used for marking the cell wall (21), in most cases it has not been clear whether they report the location(s) of cytoplasmic synthesis, periplasmic exchange, or a combination of processes. Here we show that in *M. smegmatis* the metabolic fate of monopeptide probes is partially dependent on the substituent; whereas NBD and TAMRA-conjugated d-amino acids are primarily exchanged into mycobacterial peptidoglycan by l,d-transpeptidases, their alkyne and coumarin-conjugated counterparts appear to incorporate by both extracellular and intracellular pathways (**Figure 3**). In the future, biochemical analysis of peptidoglycan composition will allow better quantitation of probe incorporation via different uptake pathways (39).

We show that dipeptide probes rescue the growth of a DdlA mutant (**Figure 4B**) and incorporate into lipid-linked peptidoglycan precursors (**Figures 5B, 5C**). To our knowledge, this is the first direct demonstration that peptidoglycan precursors can be metabolically labeled *in vivo* without radioactivity. Labeling by alkDADA unexpectedly decreased in the absence of l,d-transpeptidases (**Figure 4—figure supplement 2**). Unlike the monopeptide probes, however, alkDADA-derived fluorescence was stable to pre-treatment with imipenem, an antibiotic that targets this class of enzymes (**Figure 3**, (42)). These data suggest that the dipeptide is unlikely to be a direct substrate for l,d-transpeptidases. It is possible that d,d-carboxypeptidases cleave a small proportion of alkDADA prior to incorporation and release d-alanine and alkDA. In this scenario, apparent alkDADA labeling in *M. smegmatis* may be a combination of intracellular alkDADA incorporation via MurF (**Figures 4, 5**), extracellular alkDA incorporation by l,d-transpeptidases (**Figure 3**), and more limited intracellular alkDA incorporation by DdlA/MurF (**Figure 3**). This model is consistent with the promiscuous targeting of mycobacterial d,d-carboxypeptidases by carbapenems (42) and the ~2000-fold more efficient incorporation of alkDA compared to alkDADA (**Figure 4—figure supplement 1**). Alternatively or additionally, alkDADA may be degraded abiotically or by other enzymes such as l,d-carboxypeptidases. Finally, loss of l,d-transpeptidases may introduce global alterations in peptidoglycan metabolism. Importantly, despite differences in overall fluorescence between dipeptide-labeled wildtype and *ΔldtABE* (**Figure 4—figure supplement 1A**), sidewall fluorescence was preserved in the mutant (**Figure 4—figure supplement 1B**). In aggregate our data strongly suggest that the mycobacterial periphery is a site of active peptidoglycan synthesis.

The lateral surface of mycobacteria does not appear to contribute to cell elongation under normal growth conditions (**Figure 5—figure supplement 4**, (3–5, 7, 14)) but nevertheless hosts a substantial portion of envelope synthesis and remodeling. The intracellular difference in signal between the poles reflected relative elongation rates, as the RADA-bright cell tip, which coincides with the alkDADA- and OalkTMM-bright cell tip (**Figure 2B**) grows faster than the RADA-dim cell tip (**Figure 5—figure supplement 4**). The 2-3-fold ratio of fast/bright:slow/dim pole fluorescence roughly corresponds to previous estimates of intracellular differences in polar elongation (3, 9). Compared to *M. smegmatis*, the distribution of HADA labeling in *M. tuberculosis* is shifted away from the fast pole towards the periphery (**Figure 2D**). Diminished polarity and asymmetry is also apparent in the sub-population of *M. tuberculosis* that is labeled by alkDADA and OalkTMM (**Figure 2—figure supplement 2C**). Our data are in agreement with the heterogeneity in polar dominance observed previously for *M. tuberculosis* (15). Although we cannot rule out a contribution from cyan autofluorescence, these experiments also suggest that sidewall envelope metabolism may be even more prominent in *M. tuberculosis* than in *M. smegmatis*, comprising half of the total cell output.

What is the physiological role for cell wall synthesis that does not directly contribute to growth or division? It is possible that peptidoglycan assembly along the lateral surface of the mycobacterial cell is simply a byproduct of synthetic enzymes that are en route to the polar elongasome or the divisome. Having active enzymes at the ready could enable efficient coordination between cell growth and septation. We think that this model is less likely, however, given the energetic cost of producing complex macromolecules and the known limits on the steady-state pools of lipid-linked peptidoglycan precursors (65). Instead we propose that cell wall synthesis along the periphery could allow mycobacteria to edit what would otherwise be an inert surface (**Figure 6A**). Peptidoglycan and mycomembrane metabolism in this region may thicken or fill in the gaps of envelope that was initially deposited at the poles or the septum and enable the bacterium to correct stochastic defects and repair damage. In support of this model, we find that cell wall synthesis along the sidewall is enhanced upon exposure to peptidoglycan-degrading enzymes (**Figure 6B**). More broadly, the ability to tailor the entire cell surface, not just the ends, should enable rapid adaptation to external stimuli. Such activity may be particularly important for *M. tuberculosis*, a slow-growing organism that must survive a hostile, nutrient-poor environment.

## Materials and Methods

### Key Resources Table

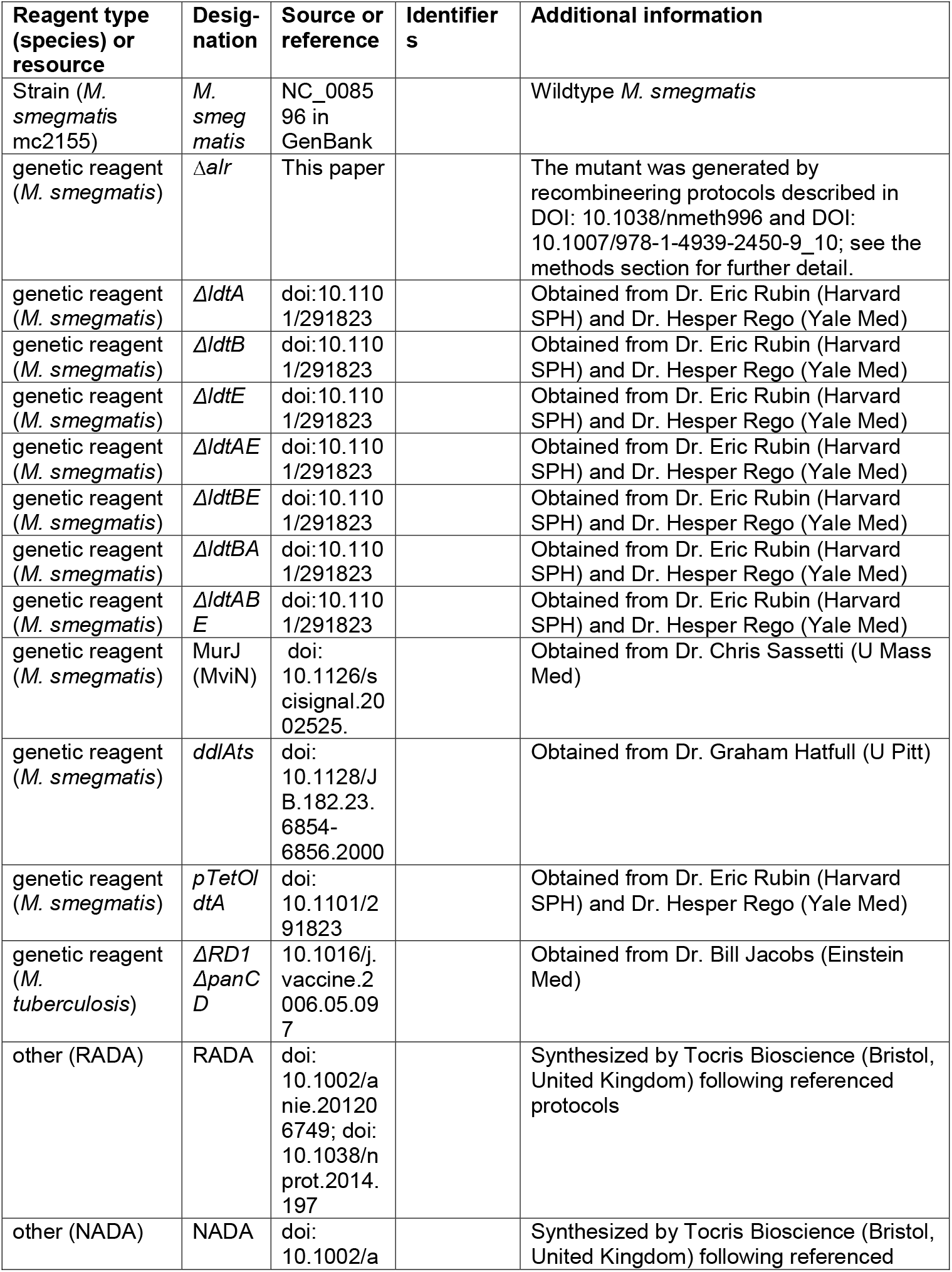
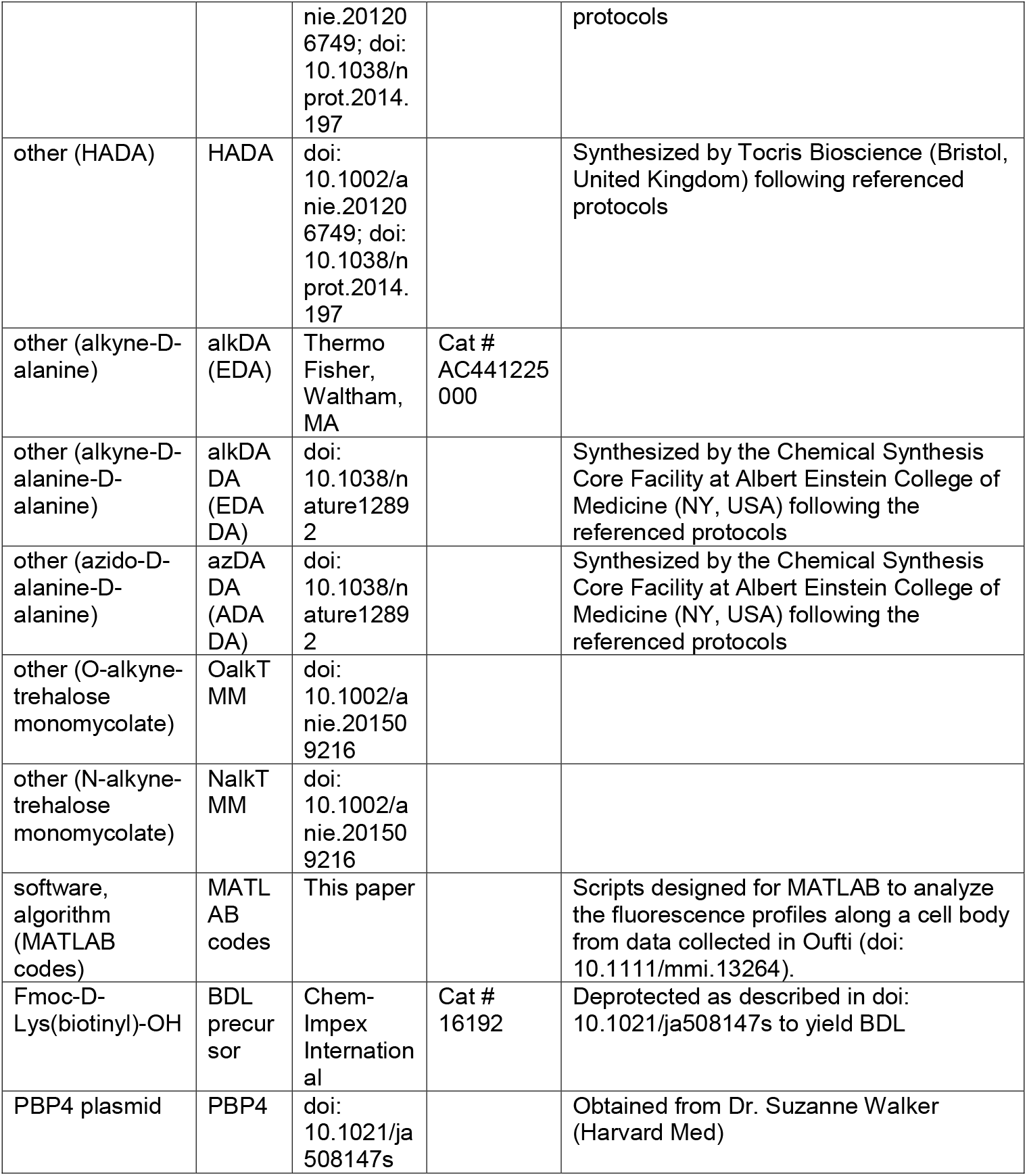

### Bacterial strains and culture conditions

mc^2^155 *M. smegmatis* and *ΔRD1 ΔpanCD M. tuberculosis* (66) were grown at 37 °C in Middlebrook 7H9 growth medium (BD Difco, Franklin Lakes, NJ) supplemented with glycerol, Tween 80 and ADC (*M. smegmatis*) or OADC and 50 μg/ml pantothenic acid (*M. tuberculosis*). The *Δalr*strain was further supplemented with 1 mM d-alanine. The *ddlA^ts^* strain was grown in 7H9-ADC at 30 °C or 37 °C as specified.

### Cell envelope labeling

Probes used in this study include (i) fluorescent d-amino acids HADA, NADA and RADA (Tocris Bioscience, Bristol, United Kingdom) (ii) alkDA (R-propargylglycine, Thermo Fisher, Waltham, MA) (iii) alkDADA and azDADA (Chemical Synthesis Core Facility, Albert Einstein College of Medicine, Bronx, NY) and (iv) OalkTMM and NalkTMM, synthesized as described previously (36). The labeling procedures were performed with modifications from (11, 13, 36). Unless otherwise indicated, mid-log *M. smegmatis* or *M. tuberculosis* were labeled with 500 μM HADA, 25 μM NADA or RADA, 50 μM alkDA, 1 or 2 mM alkDADA, 50 μM OalkTMM or 250 μM NalkTMM for 15 min or 2 hours, respectively. In some cases *M. smegmatis* was preincubated +/-antibiotics at different fold-MIC (MICs = 80 μg/mL d-cycloserine, 8 μg/mL ampicillin (with 5 μg/mL clavulanate), 0.5 μg/mL imipenem (with 5 μg/mL clavulanate), 6 μg/mL vancomycin) or +/-500 μg/mL lysozyme and 500 U/mL mutanolysin for 30 min then grown for an additional 15 min in the presence of the probes. Cultures were then centrifuged at 4 °C, washed in pre-chilled PBS containing 0.05% Tween 80 and 0.01% BSA (PBSTB) and fixed for 10 min in 2% formaldehyde at room temperature (RT). After two washes in PBSTB, the bacterial pellets were resuspended in half of the original volume of freshly-prepared CuAAC solution in PBSTB (13) containing either AF488 picolyl azide or carboxyrhodamine 110 (CR110) azide (Click Chemistry Tools, Scottsdale, AZ) or 3-azido-7-hydroxycoumarin (Jena Biosciences, Jena, Germany). For *M. tuberculosis* experiments and in **Figure 2—figure supplement 1A**, we used a modified, low-copper CuAAC reaction protocol: 200 μM CuSO_4_ and 800 μM BTTP (Chemical Synthesis Core Facility, Albert Einstein College of Medicine, Bronx, NY) were pre-mixed then added to PBSTB. Immediately before resuspending the pellets, 2.5 mM freshly prepared sodium ascorbate and 300 μM 3-azido-7-hydroxycoumarin were added to the mixture. After 30-60 min gentle agitation at RT, cultures were washed once with PBSTB, once with PBS and resuspended in PBS for imaging or flow cytometry analysis (FITC, BV510, and Texas Red channels on a BD DUAL LSRFortessa, UMass Amherst Flow Cytometry Core Facility). NHS ester dye labeling and fluorescent vancomycin labeling were performed as described (3, 6). *M. tuberculosis* was post-fixed with 4% formaldehyde overnight at RT prior to removing from the biosafety cabinet. For Bocillin labeling experiments, 500 μL of mid-log *M. smegmatis* was washed once in PBST and resuspended in PBST containing 5 μg/mL clavulanate. Bacteria were then pre-incubated or not with 50 μg/mL ampicillin or d-cycloserine at RT with gentle agitation. After 15 min, 50 μg/mL Bocillin FL (Thermo Fisher) was added and cultures were incubated for an additional 30 min. They were then washed three times in PBST and imaged live on agar pads.

### Genetic manipulation

The *Δalr M. smegmatis* strain was generated using standard recombineering methods (67, 68). 500 bp up and downstream of *alr* were cloned on either side of the *hyg^R^* cassette flanked with *loxP* sites. After induction of *recET*, transformation and subsequent selection on hygromycin and 1 mM d-alanine, PCR was used to confirm the presence of the correct insert. Strains were then cured of the *hyg^R^* cassette by transformation with an episomal plasmid carrying the Cre recombinase and a sucrose negative selection marker. Strains were cured of this plasmid by repeated passaging in the presence of sucrose.

*M. smegmatis* lacking *ldtA, ldtB* and/or *ldtE* were generously provided by Dr. Kasia Baranowski, Dr. Eric Rubin and Dr. Hesper Rego and are described in bioRxiv https://doi.org/10.1101/291823. Briefly, strains were constructed by recombineering to replace the endogenous copies with zeocin or hygromycin resistance cassettes flanked by *loxP* sites as previously described above (14). Once the knock-outs were verified by PCR, the antibiotic resistance cassettes were removed by the expression of Cre recombinase. To complement *ΔldtABE*, a copy of *ldtA* was under the constitutive TetO promoter on a kanamycin marked vector (CT94) that integrates at the L5 phage integration site of the chromosome.

### Microscopy

Fixed bacteria were imaged either by conventional fluorescence microscopy (Nikon Eclipse E600, Nikon Eclipse Ti or Zeiss Axioscope A1 with 100x objectives) or by structured illumination microscopy (Nikon SIM-E/A1R with SR Apo TIRF 100x objective).

### Microscopy analysis

Images were processed using FIJI (69) and cells were outlined and segmented using Oufti (70). Fluorescence signals of each cell were detected using Oufti and analyzed using custom-written MATLAB codes. The fluorescence intensities that we report here have been normalized by cell area. We distinguished septating from non-septating cells using the probe fluorescence profile along the long cell axis. We used the peakfinderprogram (71) to identify peaks in the labeling profile. Because our probes label both the cell poles as well as the septum, septating cells were those that had three peaks in their labeling profile, with the middle peak positioned between 30% and 70% along the normalized long cell axis. Non-septating cells were identified as having only two peaks.

### Detection of lipid-linked peptidoglycan precursors

To detect endogenous lipid-linked peptidoglycan precursors, we adopted the assay developed in (48) with some modifications. *M. smegmatis* was inoculated in 100 mL of 7H9 medium and grown to mid-log phase at 37 °C. Where applicable, MurJ was depleted by 8 hours of anhydrotetracycline-induced protein degradation as described (50). The bacteria were then divided into 25 mL cultures that were subjected or not to freshly-prepared 80 μg/mL vancomycin and/or 10 μg/mL d-cycloserine. After one hour of incubation at 37 °C, bacteria were collected by centrifugation and cell pellets were normalized by wet weight. 200-300 mg wet pellet was resuspended in 500 μL 1% glacial acetic acid in water. 500 μL of the resuspended pellet mixture was transferred into a vial containing 500 μL of chloroform and 1 mL of methanol and kept at room temperature (RT) for 1-2 hours with occasional vortexing. The mixture was then centrifuged at 21,000x g for 10 min at RT and the supernatant was transferred into a vial containing 500 μL of 1% glacial acetic acid in water and 500 μL chloroform, and vortexed for 1 min. After centrifugation at 900x g for 2 min at RT, three phases were distinguishable: aqueous, an interface, and organic. We collected the lipids from the organic phase and from the interface and concentrated it under nitrogen. Organic extracts were resuspended in 12 μL of DMSO and then incubated with purified S. *aureus* PBP4 and biotin-d-lysine (BDL; deprotected from Fmoc-d-Lys(biotinyl)-OH; Chem-Impex International) as described (48). Upon completion of the BDL exchange reaction, 10 μL of 2X loading buffer was added to the vials. Contents were boiled at 95 °C for 5 min then run on an 18% SDS polyacrylamide gel. Biotinylated species were transferred to a PVDF membrane, blotted with streptavidin-HRP (diluted 1:10,000, Thermo-Fisher) and visualized in an ImageQuant system (GE Healthcare).

To detect lipid-linked precursors that had been metabolically labeled with alkDADA or azDADA, growth of the *Δalr* strain was initially supported by the inclusion of 2 mM d-alanine-d-alanine (Sigma-Aldrich) in the 7H9 medium. *Δalr* bacteria were harvested and washed twice in sterile PBST prior to resuspension in 100 mL of pre-warmed medium. Both wildtype and *Δalr M. smegmatis* were incubated in 0.5-1 mM of alkDADA or azDADA for 1 hour then harvested as described above. Organic extracts from metabolically-labeled cultures were subjected to CuAAC reaction by adding, in order and in a non-stick vial: 2 μL of PBST, 1 μL of 5 mM CuSO_4_, 1 μL of 20 mM BTTP, 1 μL of freshly-prepared 50 mM sodium ascorbate, 3 μL of 10 mM picolyl azide biotin or alkyne biotin (Click Chemistry Tools), and 2 μL of the organic extract. The reaction was incubated for one hour at RT with gentle shaking prior to detecting as above.

## Acknowledgments

We are grateful to Dr. Kasia Baranowski, Dr. Eric Rubin and Dr. Hesper Rego for sharing l,d-transpeptidase mutants and for providing critical feedback. We thank Dr. Suzanne Walker and Dr. Kaitlin Schaefer for the S. *aureus pbp4* expression construct and for helpful technical advice; Dr. Steven Sandler, Dr. Peter Chien, Dr. James Chambers and Dr. Amy Burnside for microscopy and flow cytometry guidance; Ms. Sylvia Rivera for technical assistance; Dr. Krista Gile for guidance on statistical analysis. We also acknowledge Dr. Chris Sassetti for the MurJ depletion strain, Dr. Graham Hatfull for the *ddlA^ts^* mutant, Dr. William Jacobs for *ΔRD1 ΔpanCD M. tuberculosis* and Dr. Yves Brun, Dr. Erkin Kuru and Dr. Michael VanNieuwenhze for the initial supply of the alkDADA (EDA-DA) probe.

## Supplemental Figure Legends

**Figure 2—figure supplement 1. Metabolic labeling by azDADA comprises polar gradients in live *M. smegmatis*.**

Bacteria were labeled with azDADA for 15 min and subjected to either A, low copper CuAAC (with BTTP ligand), or B, SPAAC. Longer arrow, polar labeling. Short arrow, sidewall labeling.

**Figure 2—figure supplement 2. Heterogeneous envelope probe labeling in *M. tuberculosis*.**

A, *M. tuberculosis* was incubated for 2 hrs in NADA (green), HADA (blue) or RADA (red), washed and fixed. Fluorescence was assessed by flow cytometry. Note the cyan autofluorescence in middle panel.

B, *M. tuberculosis* was labeled with an amine-reactive dye (NHS488, black) then incubated or not for 2 hrs with HADA (dark blue), alkDA, alkDADA or OalkTMM. Alkynyl probes (light blue) and no probe control cells (grey) were detected by CuAAC with azido-coumarin. NHS488 labeling is heterogeneous and results in two distinct populations. While HADA labels both NHS488-positive and -negative bacteria, the two-step envelope probes preferentially incorporate into NHS488-positive *M. tuberculosis*.

C, Fluorescence microscopy of HADA, alkDADA and OalkTMM-labeled bacteria in B. Envelope labeling is apparent both along the sidewall, where it colocalizes with NHS488, as well as at the cell poles, where it does not. Dotted white lines highlight cell contours.

**Figure 3—figure supplement 1. Schematic of peptidoglycan synthesis in mycobacteria**.

TPase, transpeptidase.

**Figure 3—figure supplement 2. Antibiotics do not cause obvious cell death in 45 min**.

*M. smegmatis* cultures were treated with 2X MIC of indicated antibiotics for 45 min, washed, and 10-fold serial dilutions were spotted onto LB agar.

**Figure 4—figure supplement 1. alkDA labels at much lower concentrations than alkDADA**.

*M. smegmatis* were labeled with 2 mM of alkDADA (dark grey) or different concentrations of alkDA (light grey) for 15 min then fixed and subjected to CuAAC. MFI, median fluorescence intensity from which the control (no probe but subjected to CuAAC) was subtracted. Error bars, +/- standard deviation.

**Figure 4—figure supplement 2. Loss of LdtA, LdtB and/or LdtE decreases both RADA and alkDADA labeling**.

A, Wildtype and *ldt* strains were labeled with RADA (red) or alkDADA (grey) for 30 min, then washed and fixed. alkDADA detected by CuAAC with azido-CR 110. Fluorescence was quantitated by flow cytometry. Data are expressed as percentage of untreated wildtype labeling as in **Figure 3B**. Experiment was performed 4-11 times in triplicate. Error bars, +/- standard deviation. Differences within RADA- and alkDADA-labeled strains are significant at p < 0.005, one-way ANOVA comparison of log_10_ data for biological replicates.

B, Wildtype (dark green) and *ΔldtABE*(light green) *M. smegmatis* were labeled with alkDADA for 15 min, then washed, fixed and subjected to CuAAC with azido-CR110 prior to imaging. Fluorescence was quantitated for 40<n<70. Signal was normalized by cell length and brighter poles oriented to the right-hand side of the graph. AU, arbitrary units. Experiment repeated twice with similar results. Representative images at right.

**Figure 4—figure supplement 3. Loss of LdtA, LdtB and LdtE do not cause a growth defect in *M. smegmatis*.**

Culture turbidity increases at the same rate for wildtype and *ΔldtABE* growing in 7H9 medium.

**Figure 5—figure supplement 1. Short incubation in alkDADA results in polar and sidewall labeling**.

Structured illumination microscopy of *M. smegmatis* incubated in alkDADA for 2 min then washed, fixed and subjected to CuAAC with azido-CR110. Arrows highlight sidewall signal.

**Figure 5—figure supplement 2. Fluorescent vancomycin (vanc-fl) labeling at poles and sidewall**.

*M. smegmatis* were incubated in the fluorescent antibiotic for 3 hrs or ~one doubling. Longer arrow, polar labeling. Short arrow, sidewall labeling.

**Figure 5—figure supplement 3. Penicillin-binding proteins are present along mycobacterial cell periphery**.

A, *M. smegmatis* merodiploid expressing PonA1-mRFP imaged by structured illumination microscopy. Arrows highlight sidewall signal.

B, Bocillin labeling of live *M. smegmatis* pre-treated or not with ampicillin or d-cycloserine. No Boc, autofluorescence; no abx, no antibiotic.

C, Quantitation of cellular fluorescence for B. 60<n<180. Signal was normalized by cell length and brighter poles oriented to the right-hand side of the graph. AU, arbitrary units.

**Figure 5—figure supplement 4. Physical expansion of the mycobacterial cell is confined to the poles and occurs more rapidly at the RADA-bright tip**.

*M. smegmatis* were incubated in RADA for 15 min, washed and grown for 0 or 15 min in the absence of probe. Images obtained by conventional fluorescence microscopy, left, were quantified for 24<n<41 cells, right, as in **Figure 5—figure supplement 3C**. Magnification of the dim and bright pole quantitation are shown above the main graph. Distances between local maxima are expressed as percentage of total, normalized cell length. Experiment was performed twice with similar results. dp, dim pole; sw, sidewall; bp, bright pole. AU, arbitrary units.

## References

1. de Pedro MA, Quintela JC, Holtje JV, Schwarz H. 1997. Murein segregation in *Escherichia coli*. Journal of Bacteriology 179:2823–2834.

2. Daniel RA, Errington J. 2003. Control of cell morphogenesis in bacteria: two distinct ways to make a rod-shaped cell. Cell 113:767–776.

3. Aldridge BB, Fernandez-Suarez M, Heller D, Ambravaneswaran V, Irimia D, Toner M, Fortune SM. 2012. Asymmetry and aging of mycobacterial cells lead to variable growth and antibiotic susceptibility. Science 335:100–104.

4. Santi I, Dhar N, Bousbaine D, Wakamoto Y, McKinney JD. 2013. Single-cell dynamics of the chromosome replication and cell division cycles in mycobacteria. Nature Communications 4:2470.

5. Meniche X, Otten R, Siegrist MS, Baer CE, Murphy KC, Bertozzi CR, Sassetti CM. 2014. Subpolar addition of new cell wall is directed by DivIVA in mycobacteria. PNAS.

6. Thanky NR, Young DB, Robertson BD. 2007. Unusual features of the cell cycle in mycobacteria: polar-restricted growth and the snapping-model of cell division. Tuberculosis 87:231–236.

7. Kieser KJ, Rubin EJ. 2014. How sisters grow apart: mycobacterial growth and division. Nature Reviews Microbiology 12:550–562.

8. Singh B, Nitharwal RG, Ramesh M, Pettersson BM, Kirsebom LA, Dasgupta S. 2013. Asymmetric growth and division in Mycobacterium spp.: compensatory mechanisms for non-medial septa. Molecular Microbiology 88:64–76.

9. Joyce G, Williams KJ, Robb M, Noens E, Tizzano B, Shahrezaei V, Robertson BD. 2012. Cell division site placement and asymmetric growth in mycobacteria. PLoS ONE 7:e44582.

10. Brown PJ, de Pedro MA, Kysela DT, Van der Henst C, Kim J, De Bolle X, Fuqua C, Brun YV. 2012. Polar growth in the Alphaproteobacterial order Rhizobiales. PNAS 109:1697–1701.

11. Kuru E, Hughes HV, Brown PJ, Hall E, Tekkam S, Cava F, de Pedro MA, Brun YV, VanNieuwenhze MS. 2012. In Situ probing of newly synthesized peptidoglycan in live bacteria with fluorescent D-amino acids. Angewandte Chemie 51:12519–12523.

12. Zupan JR, Cameron TA, Anderson-Furgeson J, Zambryski PC. 2013. Dynamic FtsA and FtsZ localization and outer membrane alterations during polar growth and cell division in *Agrobacterium tumefaciens*. PNAS 110:9060–9065.

13. Siegrist MS, Whiteside S, Jewett JC, Aditham A, Cava F, Bertozzi CR. 2013. (D)-amino acid chemical reporters reveal peptidoglycan dynamics of an intracellular pathogen. ACS Chemical Biology 8:500–505.

14. Boutte CC, Baer CE, Papavinasasundaram K, Liu W, Chase MR, Meniche X, Fortune SM, Sassetti CM, Ioerger TR, Rubin EJ. 2016. A cytoplasmic peptidoglycan amidase homologue controls mycobacterial cell wall synthesis. eLife 5.

15. Botella H, Yang G, Ouerfelli O, Ehrt S, Nathan CF, Vaubourgeix J. 2017. Distinct Spatiotemporal Dynamics of Peptidoglycan Synthesis between *Mycobacterium smegmatis* and *Mycobacterium tuberculosis*. mBio 8.

16. Schubert K, Sieger B, Meyer F, Giacomelli G, Bohm K, Rieblinger A, Lindenthal L, Sachs N, Wanner G, Bramkamp M. 2017. The Antituberculosis Drug Ethambutol Selectively Blocks Apical Growth in CMN Group Bacteria. mBio 8.

17. Rodriguez-Rivera FP, Zhou X, Theriot JA, Bertozzi CR. 2018. Acute modulation of mycobacterial cell envelope biogenesis by front-line TB drugs. Angewandte Chemie.

18. Hett EC, Chao MC, Rubin EJ. 2010. Interaction and modulation of two antagonistic cell wall enzymes of mycobacteria. PLoS Pathogens 6:e1001020.

19. Kieser KJ, Boutte CC, Kester JC, Baer CE, Barczak AK, Meniche X, Chao MC, Rego EH, Sassetti CM, Fortune SM, Rubin EJ. 2015. Phosphorylation of the Peptidoglycan Synthase PonA1 Governs the Rate of Polar Elongation in Mycobacteria. PLoS Pathogens 11:e1005010.

20. Plocinski P, Ziolkiewicz M, Kiran M, Vadrevu SI, Nguyen HB, Hugonnet J, Veckerle C, Arthur M, Dziadek J, Cross TA, Madiraju M, Rajagopalan M. 2011. Characterization of CrgA, a new partner of the *Mycobacterium tuberculosis* peptidoglycan polymerization complexes. Journal of Bacteriology 193:3246–3256.

21. Siegrist MS, Swarts BM, Fox DM, Lim SA, Bertozzi CR. 2015. Illumination of growth, division and secretion by metabolic labeling of the bacterial cell surface. FEMS Microbiology Reviews 39:184–202.

22. de Pedro MA, Cava F. 2015. Structural constraints and dynamics of bacterial cell wall architecture. Frontiers in Microbiology 6:449.

23. Glauner B, Holtje JV. 1990. Growth pattern of the murein sacculus of *Escherichia coli*. The Journal of Biological Chemistry 265:18988–18996.

24. Liechti GW, Kuru E, Hall E, Kalinda A, Brun YV, VanNieuwenhze M, Maurelli AT. 2014. A new metabolic cell-wall labelling method reveals peptidoglycan in *Chlamydia trachomatis*. Nature 506:507–510.

25. Sarkar S, Libby EA, Pidgeon SE, Dworkin J, Pires MM. 2016. In Vivo Probe of Lipid II-Interacting Proteins. Angewandte Chemie 55:8401–8404.

26. Boyce M, Carrico IS, Ganguli AS, Yu SH, Hangauer MJ, Hubbard SC, Kohler JJ, Bertozzi CR. 2011. Metabolic cross-talk allows labeling of O-linked beta-N-acetylglucosamine-modified proteins via the N-acetylgalactosamine salvage pathway. PNAS 108:3141–3146.

27. Qin W, Qin K, Fan X, Peng L, Hong W, Zhu Y, Lv P, Du Y, Huang R, Han M, Cheng B, Liu Y, Zhou W, Wang C, Chen X. 2017. Artificial Cysteine S-Glycosylation Induced by Per-O-Acetylated Unnatural Monosacharides during Metabolic Glycan Labeling. Angewandte Chemie.

28. Mahal LK, Yarema KJ, Bertozzi CR. 1997. Engineering chemical reactivity on cell surfaces through oligosaccharide biosynthesis. Science 276:1125–1128.

29. Yang M, Jalloh AS, Wei W, Zhao J, Wu P, Chen PR. 2014. Biocompatible click chemistry enabled compartment-specific pH measurement inside *E. coli*. Nature Communications 5:4981.

30. Puffal J, Garcia-Heredia A, Rahlwes KC, Siegrist MS, Morita YS. 2018. Spatial control of cell envelope biosynthesis in mycobacteria. Pathogens and Disease 76.

31. Hancock IC, Carman S, Besra GS, Brennan PJ, Waite E. 2002. Ligation of arabinogalactan to peptidoglycan in the cell wall of *Mycobacterium smegmatis* requires concomitant synthesis of the two wall polymers. Microbiology 148:3059–3067.

32. Hayashi JM, Luo CY, Mayfield JA, Hsu T, Fukuda T, Walfield AL, Giffen SR, Leszyk JD, Baer CE, Bennion OT, Madduri A, Shaffer SA, Aldridge BB, Sassetti CM, Sandler SJ, Kinoshita T, Moody DB, Morita YS. 2016. Spatially distinct and metabolically active membrane domain in mycobacteria. PNAS 113:5400–5405.

33. Carel C, Nukdee K, Cantaloube S, Bonne M, Diagne CT, Laval F, Daffe M, Zerbib D. 2014. *Mycobacterium tuberculosis* Proteins Involved in Mycolic Acid Synthesis and Transport Localize Dynamically to the Old Growing Pole and Septum. PLoS ONE 9:e97148.

34. Backus KM, Boshoff HI, Barry CS, Boutureira O, Patel MK, D’Hooge F, Lee SS, Via LE, Tahlan K, Barry CE, 3rd, Davis BG. 2011. Uptake of unnatural trehalose analogs as a reporter for *Mycobacterium tuberculosis*. Nature Chemical Biology 7:228–235.

35. Swarts BM, Holsclaw CM, Jewett JC, Alber M, Fox DM, Siegrist MS, Leary JA, Kalscheuer R, Bertozzi CR. 2012. Probing the mycobacterial trehalome with bioorthogonal chemistry. Journal of the American Chemical Society 134:16123–16126.

36. Foley HN, Stewart JA, Kavunja HW, Rundell SR, Swarts BM. 2016. Bioorthogonal Chemical Reporters for Selective In Situ Probing of Mycomembrane Components in Mycobacteria. Angewandte Chemie 55:2053–2057.

37. Besanceney-Webler C, Jiang H, Wang W, Baughn AD, Wu P. 2011. Metabolic labeling of fucosylated glycoproteins in Bacteroidales species. Bioorganic and Medicinal Chemistry Letters 21:4989–4992.

38. Patino S, Alamo L, Cimino M, Casart Y, Bartoli F, Garcia MJ, Salazar L. 2008. Autofluorescence of mycobacteria as a tool for detection of *Mycobacterium tuberculosis*. Journal of Clinical Microbiology 46:3296–3302.

39. Cava F, de Pedro MA, Lam H, Davis BM, Waldor MK. 2011. Distinct pathways for modification of the bacterial cell wall by non-canonical D-amino acids. The EMBO Journal 30:3442–3453.

40. Ngo JT, Adams SR, Deerinck TJ, Boassa D, Rodriguez-Rivera F, Palida SF, Bertozzi CR, Ellisman MH, Tsien RY. 2016. Click-EM for imaging metabolically tagged nonprotein biomolecules. Nature Chemical Biology 12:459–465.

41. Feng Z, Barletta RG. 2003. Roles of Mycobacterium smegmatis D-alanine:D-alanine ligase and D-alanine racemase in the mechanisms of action of and resistance to the peptidoglycan inhibitor D-cycloserine. Antimicrobial Agents and Chemotherapy 47:283–291.

42. Kumar P, Arora K, Lloyd JR, Lee IY, Nair V, Fischer E, Boshoff HI, Barry CE, 3rd. 2012. Meropenem inhibits D, D-carboxypeptidase activity in *Mycobacterium tuberculosis*. Molecular Microbiology 86:367–381.

43. Bliss CI. 1956. The calculation of microbial assays. Bacteriological reviews 20:243–258.

44. Fitzgerald JB, Schoeberl B, Nielsen UB, Sorger PK. 2006. Systems biology and combination therapy in the quest for clinical efficacy. Nature Chemical Biology 2:458–466.

45. Cordillot M, Dubee V, Triboulet S, Dubost L, Marie A, Hugonnet JE, Arthur M, Mainardi JL. 2013. In vitro cross-linking of *Mycobacterium tuberculosis* peptidoglycan by L,D-transpeptidases and inactivation of these enzymes by carbapenems. Antimicrobial Agents and Chemotherapy 57:5940–5945.

46. Barreteau H, Kovac A, Boniface A, Sova M, Gobec S, Blanot D. 2008. Cytoplasmic steps of peptidoglycan biosynthesis. FEMS Microbiology Reviews 32:168–207.

47. Belanger AE, Porter JC, Hatfull GF. 2000. Genetic analysis of peptidoglycan biosynthesis in mycobacteria: characterization of a *ddlA* mutant of *Mycobacterium smegmatis*. Journal of Bacteriology 182:6854–6856.

48. Qiao Y, Lebar MD, Schirner K, Schaefer K, Tsukamoto H, Kahne D, Walker S. 2014. Detection of lipid-linked peptidoglycan precursors by exploiting an unexpected transpeptidase reaction. Journal of the American Chemical Society 136:14678–14681.

49. Qiao Y, Srisuknimit V, Rubino F, Schaefer K, Ruiz N, Walker S, Kahne D. 2017. Lipid II overproduction allows direct assay of transpeptidase inhibition by beta-lactams. Nature Chemical Biology 13:793–798.

50. Gee CL, Papavinasasundaram KG, Blair SR, Baer CE, Falick AM, King DS, Griffin JE, Venghatakrishnan H, Zukauskas A, Wei JR, Dhiman RK, Crick DC, Rubin EJ, Sassetti CM, Alber T. 2012. A phosphorylated pseudokinase complex controls cell wall synthesis in mycobacteria. Science Signaling 5:ra7.

51. Zhao H, Patel V, Helmann JD, Dorr T. 2017. Don’t let sleeping dogmas lie: new views of peptidoglycan synthesis and its regulation. Molecular Microbiology 106:847–860.

52. Mahapatra S, Crick DC, McNeil MR, Brennan PJ. 2008. Unique structural features of the peptidoglycan of *Mycobacterium leprae*. Journal of Bacteriology 190:655–661.

53. Lavollay M, Arthur M, Fourgeaud M, Dubost L, Marie A, Veziris N, Blanot D, Gutmann L, Mainardi JL. 2008. The peptidoglycan of stationary-phase *Mycobacterium tuberculosis* predominantly contains cross-links generated by L,D-transpeptidation. Journal of Bacteriology 190:4360–4366.

54. Lavollay M, Fourgeaud M, Herrmann JL, Dubost L, Marie A, Gutmann L, Arthur M, Mainardi JL. 2011. The peptidoglycan of *Mycobacterium abscessus* is predominantly cross-linked by L,D-transpeptidases. Journal of Bacteriology 193:778–782.

55. Cho H, Wivagg CN, Kapoor M, Barry Z, Rohs PD, Suh H, Marto JA, Garner EC, Bernhardt TG. 2016. Bacterial cell wall biogenesis is mediated by SEDS and PBP polymerase families functioning semi-autonomously. Nature Microbiology:16172.

56. Meeske AJ, Riley EP, Robins WP, Uehara T, Mekalanos JJ, Kahne D, Walker S, Kruse AC, Bernhardt TG, Rudner DZ. 2016. SEDS proteins are a widespread family of bacterial cell wall polymerases. Nature 537:634–638.

57. Leclercq S, Derouaux A, Olatunji S, Fraipont C, Egan AJ, Vollmer W, Breukink E, Terrak M. 2017. Interplay between Penicillin-binding proteins and SEDS proteins promotes bacterial cell wall synthesis. Scientific Reports 7:43306.

58. Arora D, Chawla Y, Malakar B, Singh A, Nandicoori VK. 2018. The transpeptidase PbpA and noncanonical transglycosylase RodA of *Mycobacterium tuberculosis* play important roles in regulating bacterial cell lengths. The Journal of Biological Chemistry 293:6497–6516.

59. Kieser KJ, Baranowski C, Chao MC, Long JE, Sassetti CM, Waldor MK, Sacchettini JC, loerger TR, Rubin EJ. 2015. Peptidoglycan synthesis in *Mycobacterium tuberculosis* is organized into networks with varying drug susceptibility. PNAS 112:13087–13092.

60. Landgraf D, Okumus B, Chien P, Baker TA, Paulsson J. 2012. Segregation of molecules at cell division reveals native protein localization. Nature Methods 9:480–482.

61. Kocaoglu O, Carlson EE. 2013. Penicillin-binding protein imaging probes. Current Protocols in Chemical Biology 5:239–250.

62. Rego EH, Audette RE, Rubin EJ. 2017. Deletion of a mycobacterial divisome factor collapses single-cell phenotypic heterogeneity. Nature 546:153–157.

63. Kanetsuna F. 1980. Effect of lysozyme on mycobacteria. Microbiology and immunology 24:1151–1162.

64. Melzer E.S.; Chambers, J.J.; Siegrist, M.S. Accepted. DivlVA concentrates mycobacterial cell envelope assembly for initiation and stabilization of polar growth. Cytoskeleton.

65. van Heijenoort Y, Gomez M, Derrien M, Ayala J, van Heijenoort J. 1992. Membrane intermediates in the peptidoglycan metabolism of *Escherichia coli:* possible roles of PBP 1b and PBP 3. Journal of Bacteriology 174:3549–3557.

66. Sambandamurthy VK, Derrick SC, Hsu T, Chen B, Larsen MH, Jalapathy KV, Chen M, Kim J, Porcelli SA, Chan J, Morris SL, Jacobs WR, Jr. 2006. *Mycobacterium tuberculosis* DeltaRD1 DeltapanCD: a safe and limited replicating mutant strain that protects immunocompetent and immunocompromised mice against experimental tuberculosis. Vaccine 24:6309–6320.

67. van Kessel JC, Hatfull GF. 2007. Recombineering in *Mycobacterium tuberculosis*. Nature Methods 4:147–152.

68. Murphy KC, Papavinasasundaram K, Sassetti CM. 2015. Mycobacterial recombineering. Methods in Molecular Biology 1285:177–199.

69. Schindelin J, Arganda-Carreras I, Frise E, Kaynig V, Longair M, Pietzsch T, Preibisch S, Rueden C, Saalfeld S, Schmid B, Tinevez JY, White DJ, Hartenstein V, Eliceiri K, Tomancak P, Cardona A. 2012. Fiji: an open-source platform for biological-image analysis. Nature Methods 9:676–682.

70. Paintdakhi A, Parry B, Campos M, Irnov I, Elf J, Surovtsev I, Jacobs-Wagner C. 2016. Oufti: an integrated software package for high-accuracy, high-throughput quantitative microscopy analysis. Molecular Microbiology 99:767–777.

71. Sliusarenko O, Heinritz J, Emonet T, Jacobs-Wagner C. 2011. High-throughput, subpixel precision analysis of bacterial morphogenesis and intracellular spatio-temporal dynamics. Molecular Microbiology 80:612–627.

